# Evaluating polarizable biomembrane simulations against experiments

**DOI:** 10.1101/2023.12.01.569661

**Authors:** Hanne S. Antila, Sneha Dixit, Batuhan Kav, Jesper J. Madsen, Markus S. Miettinen, O. H. Samuli Ollila

## Abstract

Owing to the increase of available computational capabilities and the potential for providing more accurate description, polarizable molecular dynamics force fields are gaining popularity in modelling biomolecular systems. It is, however, crucial to evaluate how much precision is truly gained with the increased cost and complexity of the simulation. Here, we leverage the NMRlipids open collaboration and Databank to assess the performance of available polarizable lipid models—the CHARMM-Drude and the AMOEBA-based parameters—against high-fidelity experimental data and compare them to the top-performing non-polarizable models. While some improvement in the description of ion binding to membranes is observed in the most recent CHARMM-Drude parameters, and the conformational dynamics of AMOEBA-based parameters are excellent, the best non-polarizable models tend to outperform their polarizable counterparts for each property we explored. The identified shortcomings range from inaccuracies in describing the conformational space of lipids to excessively slow conformational dynamics. Our results provide valuable insights for further refinement of polarizable lipid force fields and for selecting the best simulation parameters for specific applications.

## 1 Introduction

Classical molecular dynamics (MD) simulations are nowadays widely and almost routinely used to model a wide range of biomolecular complexes. ^1^ In conventional MD models (known as force fields), the electrostatic interactions are described by assigning the atoms and molecules with static point charges. Dynamic effects arising from electronic polarizability are thus not explicitly included, but only considered in an averaged fashion within the force field parameterization process, where parameters are obtained by fitting to macroscopic observables or to *ab initio* calculations. However, electronic polarization is perceived to be a key contribution to correctly describe many biomolecular systems—including water, ion hydration and ion binding to molecules, cation–*π* and *π*–*π* interactions,^2^ the vibrational Stark effect,^3,4^ as well as co-operativity in interactions in general. ^5^ These low-level interactions also translate to the behaviour of large-scale biomolecular systems, such as ion channels where ion-selectivity and ion currents may be affected by polarization, ^6–9^ and telomeric DNA^10^ where the conformations adopted are mediated by ionic interactions. Consequently, significant efforts have been dedicated to introduce explicit polarizability into MD simulations in the hopes of reaching a more accurate representation of reality.^11–18^

In a bilayer membrane, specifically, the molecular (dielectric) environment varies dramatically when crossing from the water phase across the dipolar/charged lipid headgroup interface to the hydrophobic tail region. Therefore, including polarizability in the lipids is expected to improve the membrane potential and especially the description of membrane binding processes, of the translocation of charged biomolecules across the membrane, and of the behavior of molecules residing within membranes, such as membrane proteins.^16,17,19–25^ However, the quality of polarizable lipid models has not been evaluated on an equal footing with the non-polarizable models.

The currently available lipid force fields with explicit electronic polarization include the CHARMM-Drude,^19,26^ AMOEBA-based,^20,27^ and CHARMM-Fluctuating Charge (FQ)^28^ parameters. Their underlying strategies differ: 1) the classical Drude oscillator (CHARMM-Drude) models polarization by two separate (core and shell) charges connected with a spring that orients and stretches in response to the environment, giving the site a fluctuating dipole moment;^16^ 2) the induced point dipole/multipole approach of AMOEBA (Atomic Multipole Optimized Energetics for Biomolecular Applications) uses polarizable point dipoles placed on chosen sites of the molecule;^29^ and 3) the electronegativity equilization (fluctuating charge, FQ) employs atomic charges that are not constant but can redistribute within the molecule according to the electronegativities of the molecule atoms and the electric fields from their molecular environment.^30^ All of these approaches result in an increasing computational cost, e.g, by introducing new types of interactions, more interactions sites, or by requiring a shorter time step. As a computationally efficient alternative approach, the electronic continuum correction (ECC) has been proposed to implicitly include polarizability by scaling the atom partial charges. ^23,31^

Our previous efforts in benchmarking state-of-the-art non-polarizable lipid force fields have demonstrated that the quality of predictions for important membrane properties greatly varies between different force fields, particularly for lipid headgroup conformational ensembles and ion binding affinities.32–40 While the ability to capture these membrane properties correctly is important on its own right, it also creates the basis for the description of more complex systems: For example, ion binding affinity regulates membrane surface charge, and having a wide variety of conformations available for lipid headgroups appears essential for capturing realistic protein–lipid interactions.35 Consequently, such benchmark studies are urgently needed also for polarizable lipid force fields, in particular considering the increased computational cost they come with and their pledge to capture a broader range of physical phenomena at the polar membrane regions.

Here we assess the quality of the currently actively developed polarizable lipid force fields, the CHARMM-Drude^19,26^ and AMOEBA-based^20,27^ parameters, using the resources and framework of the NMRlipids open collaboration (nmrlipids.blogspot.fi). These two force fields were selected for comparison, because they are increasingly used in biomolecular simulations and have parameters available for several lipids for which the corresponding experimental data are available in the NMRlipids Databank (databank.nmrlipids.fi).^40^ We assess the structural quality of POPC (1-palmitoyl-2-oleoyl-sn-glycero-3-phosphocholine), DOPC (1,2-dioleoyl-sn-glycero-3-phosphocholine), and POPE (1-palmitoyl-2-oleoyl-sn-glycero-3-phosphoethanolamine) lipid bilayer simulations against experimental Nuclear Magnetic Resonance (NMR) spectroscopy and Small Angle X-ray Scattering (SAXS) data using the quality metrics defined in the NMRlipids Databank.^40^ Cation binding to membranes is evaluated against salt-induced changes in the NMR C–H bond order parameters, ^33^ and lipid headgroup conformational dynamics are benchmarked to data from NMR spin relaxation rate experiments.^36,41^ Furthermore, for each experimental benchmark we compare the polarizable models to the best-performing non-polarizable simulations in the NMRlipids Databank.^40^ Our results will act as a useful reference for selecting the best polarizable lipid models for a wide range of applications and as a guide for future development of polarizable force field parameters for lipids.

## 2 Methods

### 2.1 Using a polarizable force field for membrane simulations

While non-polarizable MD simulations of membranes can nowadays be routinely performed with several simulation engines and force fields, polarizable simulations still bear many practical complications. Out of the currently popular MD simulation packages for membrane simulations, OpenMM^42^ supports both AMOEBA and Drude force fields, NAMD^43^ can only run the Drude force field, whereas GROMACS^44^ has only limited support for the Drude polarizable force field via an unofficial Git-branch.^45^ TINKER^46^ is widely used with AMOEBA, but it does not support the semi-isotropic pressure coupling required for membrane simulations. Consequently, we selected OpenMM for simulations in this work.

Another practical issue is the availability of parameters for the molecules of interest. For the Drude force field, CHARMM-GUI^47,48^ can generate the topology and input parameters, which greatly simplifies the employment of this model.^49^ For the AMOEBA lipid parameters, standard protocols are not available, but some parameters can be found in the literature. ^7,50^

Lastly, while conducting polarizable membrane simulations, one should consider the increased computational cost arising from the explicit treatment of electronic polarizability. For the Drude-based models, a slowdown occurs both because the addition of Drude particles increases the number of interaction pairs, and because the employed extended dual-Langevin thermostat requires a shorter 1 fs integration time step (compared to the 2 fs typically used for non-polarizable membrane simulations). The AMOEBA force field can use a multi-time-step integration algorithm, where the non-electronic interactions are iterated with a 2 fs time step and the more computationally unstable polarization terms with a shorter time step. However, the multi-time-step scheme only partly mitigates the computational cost. For the systems studied here, our AMOEBA simulations are roughly 1–2 orders of magnitude slower to run than CHARMM-Drude simulations, which in turn are *∼*4 times slower than simulating an equivalent system using non-polarizable CHARMM36.

All simulations performed in this work are listed in Table 1 with links to the openly available trajectory data. Data not mentioned in Table 1 were obtained by analysing preexisting trajectories from the NMRlipids Databank and are cited in corresponding figure captions.

**Table 1:** Systems simulated specifically for this work. Column *N_l_* gives the number of lipids, *N_w_* the number of water molecules, *T* (K) denotes the temperature in kelvins. The salt concentrations in column “Ion (M)” is calculated from the number of cations *N_c_* as [salt]=*N_c_*×[water] / *N_w_*, where [water] = 55.5 M. Simulated time is listed in column *t*_s_ and time used for analysis in *t*_a_. Column *t*_eq_ gives the relative equilibration times with respect to the trajectory lengths based on PCAlipids^76,77^ and computed using the NMRlipids Databank:^40^ *t*_eq_ *<* 1 indicates convergence, *t*_eq_ *>* 1 indicates the presence of a longer time-scale than the trajectory length. Column “files [Ref.]” gives the reference to openly accessible simulation data.

| Lipid:salt | force field | Ion (M) | $N_l$ | $N_w$ | $N_c$ | T (K) | $t_s$ (ns) | $t_a$ (ns) | $t_{\text{eq}}$ | files [Ref.] |
| --- | --- | --- | --- | --- | --- | --- | --- | --- | --- | --- |
| POPC | Drude2017 | 0 | 144 | 6400 | 0 | 303 | 500 | 400 | 5.79 | 81 |
| | Drude2023 | 0 | 72 | 2239 | 0 | 303 | 300 | 200 | $2.29 \pm 0.13$ | 82, 60 |
| POPE | Drude2017 | 0 | 144 | 6400 | 0 | 308 | 350 | 300 | 5.79 | 83 |
| | Drude2023 | 0 | 72 | 2304 | 0 | 303 | 300 | 200 | $1.71 \pm 0.10$ | 59, 84 |
|  | AMOEBA | 0 | 72 | 2880 | 0 | 303 | 306 | 306 | 0.43 | 85 |
| POPC:NaCl | Drude2017 | 0.350 | 144 | 6400 | 41 | 303 | 500 | 400 | 3.31 | 86 |
|  | Drude2017 | 0.450 | 144 | 6400 | 51 | 303 | 500 | 400 | 2.5 | 87 |
|  | Drude2017 | 0.650 | 144 | 6400 | 77 | 303 | 500 | 400 | 3.30 | 88 |
|  | Drude2017 | 1.0 | 144 | 6400 | 115 | 303 | 500 | 400 | 4.06 | 89 |
|  | Drude2023 | 0.350 | 128 | 6400 | 41 | 303 | 224 | 220 | 2.63 | 90 |
|  | Drude2023 | 1.0 | 128 | 6400 | 115 | 303 | 220 | 220 | 2.77 | 91 |
| POPC:CaCl<sub>2</sub> | Drude2017 | 0.350 | 144 | 6400 | 41 | 303 | 500 | 400 | 2.45 | 92 |
|  | Drude2017 | 0.450 | 144 | 6400 | 52 | 303 | 500 | 400 | 2.56 | 93 |
|  | Drude2017 | 0.650 | 144 | 6400 | 76 | 303 | 500 | 400 | 4.46 | 94 |
|  | Drude2017 | 1.0 | 144 | 6400 | 114 | 303 | 500 | 400 | 5.00 | 95 |
|  | Drude2023 | 0.350 | 128 | 6400 | 41 | 303 | 219 | 219 | 2.63 | 96 |
|  | Drude2023 | 0.790 | 128 | 6400 | 91 | 303 | 214 | 214 | 4.32 | 97 |
| DOPC | AMOEBA | 0 | 72 | 2880 | 0 | 303 | 202 | 202 | 0.62 | 98 |
| DOPC:NaCl | AMOEBA | 0.450 | 72 | 2880 | 17 | 303 | 218 | 218 | 0.60 | 99 |
|  | AMOEBA | 1.0 | 72 | 2880 | 35 | 303 | 202 | 202 | 0.61 | 100 |
| DOPC:CaCl<sub>2</sub> | AMOEBA | 0.450 | 72 | 2880 | 16 | 303 | 218 | 218 | 0.53 | 101 |
|  | AMOEBA | 1.0 | 72 | 2880 | 36 | 303 | 218 | 218 | 0.66 | 102 |

### 2.2 Simulations with CHARMM-Drude parameters

The CHARMM-Drude2017 simulations were performed with OpenMM 7.5.0^42^ using parameters extracted with *Membrane Builder* ^51–54^ and *Drude Prepper* ^49^ from CHARMM-GUI. ^47,48^ Before starting the simulations, membrane structures were equilibrated for 200 ns using the non-polarizable CHARMM36 force field,^55^ and the last frames of these simulations were used to generate the starting structures for the polarizable force field simulations. Ion parameters were obtained from Ref. 56, and the SWM4-NDP water model^57^ was employed in all Drude simulations.

As this manuscript was prepared, the CHARMM-Drude2023 force field parameters were not integrated into CHARMM-GUI. Therefore, the simulation setups with NaCl and CaCl_2_ ^56^ using the SWM4-NDP water model^57^ were generated following the instructions in the original CHARMM-Drude2023 paper^26^ using the CHARMM program^58^ and the last frames of the 200-ns-long CHARMM36 simulations (the same ones as for CHARMM-Drude2017). The salt-free CHARMM-Drude2023 simulations were obtained from Zenodo.^59,60^

A dual Langevin thermostat was employed to keep the Drude particles at 1.0 K and the rest of the system at 303 K. A Drude-hardwall of 0.02 nm was used to keep the Drude particles close to their parent atoms. Semi-isotropic Monte Carlo barostat^61^ was used to couple pressure to 1 bar independently in the membrane plane and in the membrane normal directions. Length of the covalent bonds containing hydrogens were constrained. For CHARMM-Drude2017, Particle Mesh Ewald (PME)^62^ was used to compute the Coulomb interactions, and the van der Waals interactions were brought to zero between 1.0 nm and 1.2 nm using a switching function. For CHARMM-Drude2023 without salt, PME was used for electrostatics and the Lennard-Jones Particle Mesh Ewald (LJ-PME) method was used to compute the long-range dispersions.^63^ For CHARMM-Drude2023 with salt, same setting as for CHARMM-Drude2017 were used. Simulation frames were saved every 10 ps.

### 2.3 Simulations with AMOEBA-based parameters

For simulations with the AMOEBA-based force field, we used the OpenMM implementation of parameters developed by Chu *et al.*^27^ available on GitHub.^7,50^ All AMOEBA simulations were run using OpenMM 7.5.1.^42^ The same initial structures as in the CHARMM-Drude simulations were used. A multi-time-step Langevin integrator^64,65^ was used to iterate the bonded and non-bonded interactions with time steps of 0.5 fs and 2.0 fs, respectively. A non-bonded cutoff of 1.2 nm was applied, while semi-isotropic Monte Carlo barostat^61^ was used to couple pressure to 1 bar independently in the membrane plane and normal directions. The ion and water parameters were obtained from Ref. 66. Simulation frames were saved every 10 ps. Further simulation details can be found in the input files of the respective simulations (see links to openly available data in Table 1).

### 2.4 Choice of water model

In this study, we have used the water models that are native to the developed lipid force field: AMOEBA14^67^ for the AMOEBA force field, and SWM4-NDP^57^ for CHARMM-Drude2017 and CHARMM-Drude2023 force fields. Other water models for the AMOEBA^68–70^ and Drude^71,72^ force field families are available, and it is important to note that force fields’ predictive capabilities may be sensitive to the chosen water model^73,74^ as the force field parameters are often fine-tuned based on simulations in aqueous environment. Therefore, the results presented here are limited to the chosen water models. A complete evaluation of the effects that the choice of water model has on the dynamics and structure of polarizable lipids would be valuable to the simulation community, yet such evaluation is beyond the scope of this work.

### 2.5 Analysis of simulations

All simulations were first added to the NMRlipids Databank.^40^ Areas per lipid, SAXS form factors (*|F* (*q*)*|*), relative equilibration times t_eq_, and C–H bond order parameters (*S*_CH_) are automatically calculated by the NMRlipids Databank, ^40^ and were extracted from there. Quality evaluation metrics were quantified as detailed in the NMRlipids Databank, ^40^ with the exception that the POPC simulations at 303 K were paired with the experimental data measured at 300 K. (In the NMRlipids Databank, simulations are paired with experiments with the maximum temperature difference of two degrees). The order parameter qualities (*P* ^headgroup^, *P* ^sn^*^−^*^1^, and *P* ^sn^*^−^*^2^) reflect the average probabilities for *S*_CH_ within the corresponding molecular segment to locate within the experimentally acceptable values, taking the error bars of both simulation and experiment into account. Qualities of the SAXS form factors, FF*_q_*, depict the difference of the first *|F* (*q*)*|* minima locations in simulation and experiment; this choice avoids the effects arising from the simulation-size-dependency on *F* (*q*).^40^ Note that in quantifying the bilayer electron densities for calculation of SAXS curves, the NMR-lipids Databank analysis algorithm places electrons as point charges at the atom centers without considering the redistribution of charge density due to the polarizability. Nevertheless, we expect this approximation not to have significant effect on the resulting SAXS form factors.^75^ Relative equilibration times *t*_eq_ were calculated using the PCAlipids^76,77^ method (as implemented in the NMRlipids Databank^40^). In this analysis, each lipid configuration is first aligned to the average structure from the trajectory, and principal component analysis is then applied to the heavy atom coordinates. The distribution convergence time of the motions along the first principal component—the motions with the longest convergence time^76^—is then quantified and divided by the total trajectory length: *t*_eq_ = *t*_convergence_*/t*_s_. A relative equilibration time *t*_eq_ *>* 1 indicates that individual lipids may not have sufficiently sampled their conformational ensembles and longer simulations are advisable.

The mass density profiles were calculated using MDAnalysis^78,79^/NumPy^80^ for the CHARMM-Drude and AMOEBA simulations; and the gmx density Gromacs command for the CHARMM36 and ECClipids simulations, for which the trajectories where extracted from the NMRlipids Databank. For all calculations, the membrane center of mass was translated to the origin. All data were normalized to give probability densities of finding the particle at the given distance.

The *R*_1_ relaxation rates and the effective correlation times *τ*_eff_, along with the accompanying error estimates, were quantified from the trajectories using an in-house python script available at github.com/NMRLipids/NMRlipidsVIpolarizableFFs/tree/master/scripts/correlation as elaborated in Ref. 36.

## 3 Results and Discussion

### 3.1 Evaluation of lipid bilayer structural properties

To evaluate the structural properties of lipid bilayers in simulations with polarizable force fields, we simulated POPC and POPE lipid bilayers with CHARMM-Drude2017^19^ and CHARMM-Drude2023, ^26^ and POPE and DOPC bilayers with the AMOEBA-based^20,27^ parameters (Table 1). These systems were selected due to the simultaneous availability of both force field parameters and experimental data.^40^ We then added our simulation trajectories to the NMRlipids Databank, such that its quality metric could be used to evaluate each trajectory against experiments. ^40^ The metric measures the quality in two aspects: First, the quality of the conformational ensemble of individual lipids is evaluated against the C–H bond order parameters *S*_CH_ from NMR; and second, the consistency of membrane dimensions is compared against SAXS form factors *F* (*q*).^40,103^ The former metric is further divided into three parts that separately describe the average quality of the headgroup and glycerol backbone region (*P* ^headgroup^), and the two acyl chains (*P* ^sn^*^−^*^1^ and *P* ^sn^*^−^*^1^). While the *S*_CH_ primarily reflect lipid conformations, the acyl chain order parameters are also a good proxy for membrane packing: The smaller the area per lipid, the larger the magnitudes of the *S*_CH_ tend to be.^40^

Figure 1 shows direct comparisons between the simulated and experimental data; Table 2 shows the resulting quality metrics and comparisons to three non-polarizable force fields: OPLS3e, CHARMM36, and GROMOS-CKP. The CHARMM-Drude2017 simulations predict slightly too packed membranes (with excessively negative acyl chain C–H bond order parameters, Fig. 1) compared to experiments and to simulations with the highest quality in the NMRlipids Databank (OPLS3e for POPC and GROMOS-CKP for POPE, Table 2).^40^

**Figure 1:**
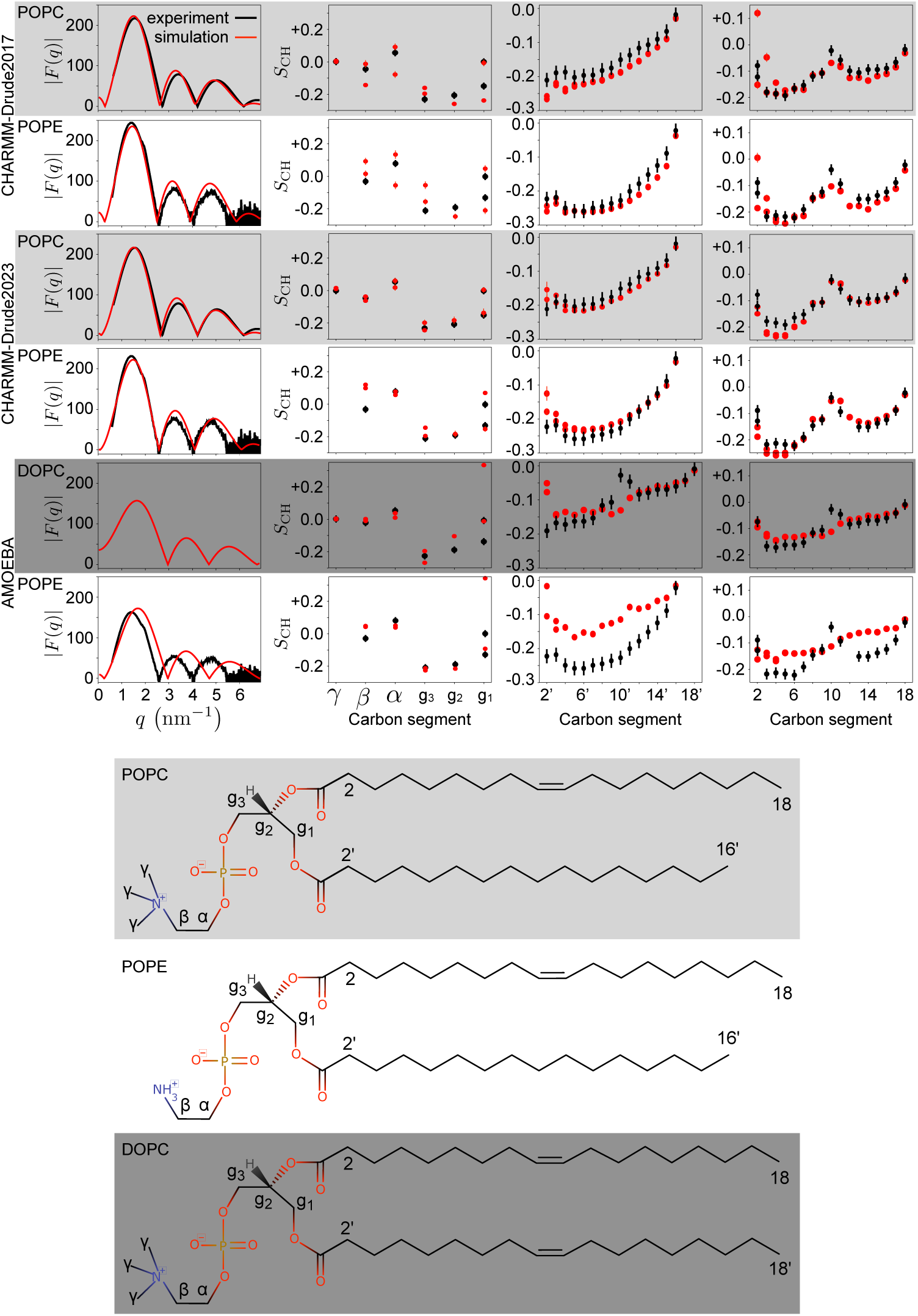
The X-ray scattering form factors *F* (*q*) (leftmost column); and the C–H bond order parameters *S*_CH_ for headgroup and glycerol backbone (second column from left), sn-1 (second column from right), and sn-2 acyl chains (rightmost column) compared between simulations (red) and experiments (black) using the NMRlipids Databank. The experimental data were originally reported in Refs. 35,40,111–113. For the CHARMM-Drude2023 simulations, we selected representative replicas among the three available ones (for all POPC replicas, see SI Fig. S1). A comparison of bilayer electron densities from which the SAXS curves are calculated is presented in SI Fig. S2. The modelled lipids and their carbon-naming scheme is shown at the bottom.

**Table 2:** NMRlipids Databank quality metrics^40^ and areas per lipid (APL) compared between simulations with polarizable force fields and the best (non-polarizable) simulations currently found in the NMRlipids Databank (OPLS3e^104^ for POPC and GROMOS-CKP^105–107^ for POPE), as well as simulations with the non-polarizable CHARMM36 force field. The segment-wise quality metrics *P* ^headgroup^, *P* ^sn^*^−^*^1^, and *P* ^sn^*^−^*^2^ reflect the average probability of the *S*_CH_ within the corresponding segment to agree with experiments (larger *P* means higher quality). The form factor quality metric, FF*_q_*, presents the difference in essential features between the simulated and experimental form factors (smaller value indicates higher quality). Experimental estimates for areas per lipid are POPC: 64.3*±*1 Å^2,108^ DOPC: 67.5*±*1 Å^2,109^ and POPE: 56.7*±*3 Å^2^.^110^

| Lipid | Force field | $P^{\text{headgroup}}$ | $P^{\text{sn}-1}$ | $P^{\text{sn}-2}$ | $\text{FF}_q$ | APL |
| --- | --- | --- | --- | --- | --- | --- |
| POPC | OPLS3e | 0.76 | 0.87 | 0.85 | 0.15 | 66.5 |
| POPC | CHARMM36 | 0.70 | 0.54 | 0.69 | 1.16 | 65.0 |
| POPC | CHARMM-Drude2017 | 0.52 | 0.29 | 0.53 | 1.06 | 62.5 |
| POPC | CHARMM-Drude2023 | 0.63 | 0.60 | 0.57 | 0.96 | 64.5 |
| POPE | GROMOS-CKP | 0.29 | 0.83 | 0.48 | 0.40 | 59.6 |
| POPE | CHARMM36 | 0.54 | 0.52 | 0.27 | 1.30 | 57.2 |
| POPE | CHARMM-Drude2017 | 0.06 | 0.53 | 0.27 | 0.80 | 56.6 |
| POPE | CHARMM-Drude2023 | 0.28 | 0.59 | 0.54 | 0.00 | 61.4 |
| POPE | AMOEBA | 0.21 | 0.10 | 0.23 | 3.80 | 66.9 |
| DOPC | AMOEBA | 0.60 | 0.60 | 0.54 | - | 70.2 |

This is similar to the non-polarizable CHARMM36 simulations. However, the quality of headgroup conformations in CHARMM-Drude2017 is worse (0.52 for POPC and only 0.06 for POPE) than in its non-polarizable counterpart (0.70 and 0.54). This is likely because the CHARMM-Drude2017 force-field parameters for the headgroup and glycerol backbone were optimized to reproduce the average absolute values of the experimentally determined *S*_CH_, that is, without taking into account the order parameter sign and “forking” (measurably different *S*_CH_ for different C–H bonds at a single carbon atom). ^37^ A better description of the PC and PE headgroups and the glycerol backbone is provided by CHARMM-Drude2023 (0.63 and 0.28), yet its quality still remains below that offered by the non-polarizable CHARMM36. Differences of headgroup conformations between force fields are shown in terms of dihedral angle distributions in Supplementary Information Fig. S3; see also discussion about the conformational dynamics in Section 3.2 below. Also the quality of membrane packing and acyl chain order are improved in CHARMM-Drude2023 (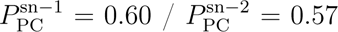 and 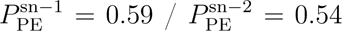 in Table 2) compared to the earlier version (0.29 / 0.53 and 0.53 / 0.27); but again, it is outperformed by the best available non-polarizable simulations (OPLS3e 0.87 / 0.85 and GROMOS-CKP 0.83 / 0.48).

The AMOEBA-based simulations capture the headgroup and glycerol backbone order parameters reasonably well—with the exception of g_1_, where forking is unacceptably large (Fig. 1). However, the experimentally observed high order parameters at the double-bond region in both DOPC acyl chains and in the sn-2 chain of POPE are not even qualitatively captured. These high order parameters signal an important mechanism through which acyl chain double bonds affect membrane properties,^114^ and are well reproduced in all the state-of-the-art non-polarizable atomistic MD force fields. ^103^ Furthermore, the AMOEBA-based parameters substantially overestimate the area per lipid in POPE simulations (Table 2), which is connected to too disordered acyl chains. Similar issues are evident also in the membrane data presented in a recent publication^115^ for the AMOEBA-based cholesterol model. The unsatisfactory description of the lipid tail region and the area per lipid is further reflected in the inability of AMOEBA-based parameters to capture the POPE SAXS curve (Fig. 1). Thus, we can conclude that the AMOEBA-based parameters used in our simulations did not reproduce essential membrane properties at the level of the state-of-the-art lipid parameters.

### 3.2 Evaluation of lipid conformational dynamics

While the C–H bond order parameters *S*_CH_ are highly sensitive to the lipid conformational ensemble, this correspondence is not unique. Essentially, the *S*_CH_ describe only the averages of the conformational distributions, furthermore they carry no information on the dynamics of the conformational sampling: A simulation that reproduces the order parameters has an ensemble that is (potentially) correct (necessary but not sufficient condition), but even the correct ensemble may not be sampled at the experimentally observed dynamics. To elucidate the dynamics of polarizable force fields, Fig. 2 compares their ^13^C NMR spin-lattice relaxation rates *R*_1_, and C–H bond effective correlation times *τ*_eff_, with experiments^41^ and the best non-polarizable simulations from our previous study.^36^ Here, we focus on the PC headgroups and glycerol backbone due to the availability of both experimental data and polarizable simulations. The *R*_1_ rates measured at typical magnetic field strengths are sensitive to rotational dynamics of C–H bonds on timescales around *∼*0.1–1 ns, while the *τ*_eff_ respond to a wide range of dynamical processes from 100 ps up to *∼*1 000 ns.^41^

**Figure 2:**
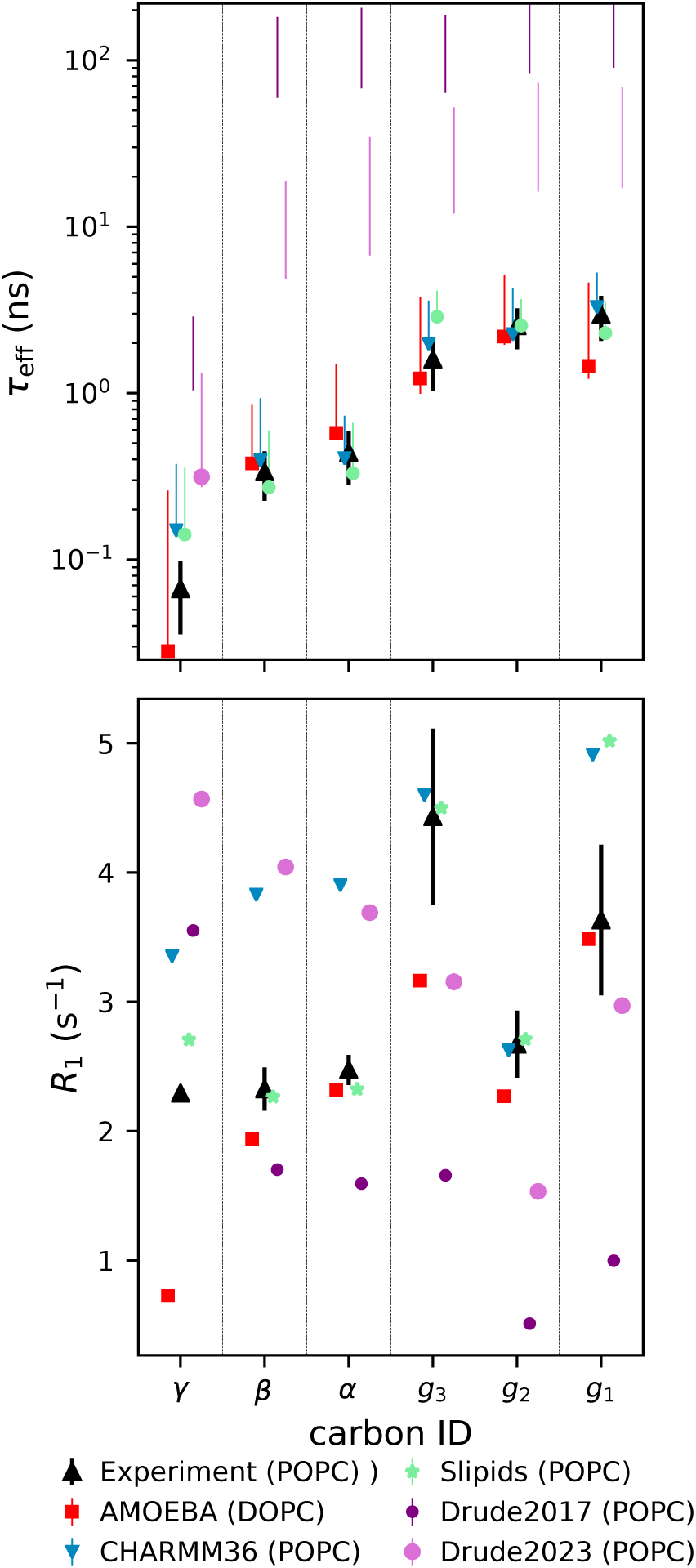
Effective correlation times *τ*_eff_ (top) and spin-lattice relaxation rates *R*_1_ (bottom) for the polarizable, and the best-performing non-polarizable (CHARMM36 and Slipids,^119^ data from Ref. 36), force fields. Note that the top panel *y*-axis is logarithmic to visualize the slow dynamics of the Drude-based models. Experimental values from Ref. 117. For the simulated *τ*_eff_, the data point quantifies the average over the C–H bonds. If *τ*_eff_ could not be determined for all bonds due to slow convergence, only the range from the mean of the lower to the mean of the upper error estimates is shown. For *R*_1_, the error bars were smaller than the symbol size. All the simulations shown here were salt-free.

The effective correlation time *τ*_eff_ gives an average measure of how fast the molecular conformations go through the phase space that leads to the average C–H bond order parameters. The *τ*_eff_ in CHARMM-Drude2017 and CHARMM-Drude2023 are approximately two and one orders of magnitude slower, respectively, than the values extracted from experiments and the best available simulations (Fig. 2). This indicates that not only are these polarizable simulations computationally costlier (due to reasons outlined in Introduction) for equivalent lengths of trajectory, but one would also have to create longer trajectories to obtain converged results. This is further evidenced by the relative equilibration times given in Table 1. By this measure, the CHARMM-Drude simulations have not converged within the rather standard trajectory lengths used in this work. The non-polarizable counterpart of the Drude models, CHARMM36, exhibits much more realistic, i.e. faster, dynamics and thus shorter *τ*_eff_ . Also the dynamics in the 1-ns range (*R*_1_ rates), are on average slightly more realistic in CHARMM36 simulations compared to both of its polarizable counterparts. The inaccuracies of the *R*_1_ rates at the glycerol region have been already pointed out upon publication of the CHARMM-Drude2023 model.^26^ Interestingly, the CHARMM-Drude models have been reported to have slower water-hydrogen-bonding dynamics around amino acids compared to their non-polarizable counterpart, ^116^ which might align with an overall slower dynamics of the model in addition to enhanced water binding.

Figure 2 also shows that, in contrast to the Drude-based models discussed above, the *R*_1_ rates and *τ*_eff_ times in DOPC simulations with the AMOEBA-based force field reproduce the experimental data from POPC well, on par with the best non-polarizable models (Slipids and CHARMM36). The small difference in acyl chain composition (DOPC vs. POPC) is not expected to affect headgroup dynamics due to the effective decoupling between the hydrophilic and hydrophobic membrane regions.^117,118^

### 3.3 Cation binding to membranes in polarizable simulations

Given the abundance of cations in biological systems, accurately capturing their interactions with membranes in simulations has the uttermost importance. A wealth of experimental evidence shows that monovalent ions (except for lithium) exhibit very weak binding affinity to PC lipid bilayers, while multivalent ions such as calcium bind more strongly.^33^ However, simulations with non-polarizable force fields (without any additional corrections) systematically overestimate cation binding to lipid bilayers.^33^ Implicit inclusion of electronic continuum correction (ECC) to both ion and lipid parameters can substantially improve the situation,^23,24,35^ suggesting that electronic polarizability plays an important role in ion binding to membranes. One might thus expect that simulations with explicitly polarizable force fields will more accurately describe ion binding to membranes. To test this notion, we evaluated ion binding to membranes using the experimental NMR “lipid electrometer” data: Here the amount of ion binding to the membrane is quantified by monitoring the change in the lipid headgroup order parameters (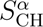 and 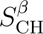) in response to an increasing salt concentration.^33,120^ Figure 3 shows the changes in these order parameters, as induced by increasing NaCl or CaCl_2_ concentration, for simulations and experiments; Fig. 4 shows the corresponding density profiles of ions with respect to the bilayer normal in simulations. Results from the AMBER-based ECClipids model are presented as a reference simulation that gives a good agreement with experiments for cation binding. ^23^ We also show data from the non-polarizable CHARMM36, where the NBFIX correction for the ion models was specifically developed to address overbinding.^121,122^

**Figure 3:**
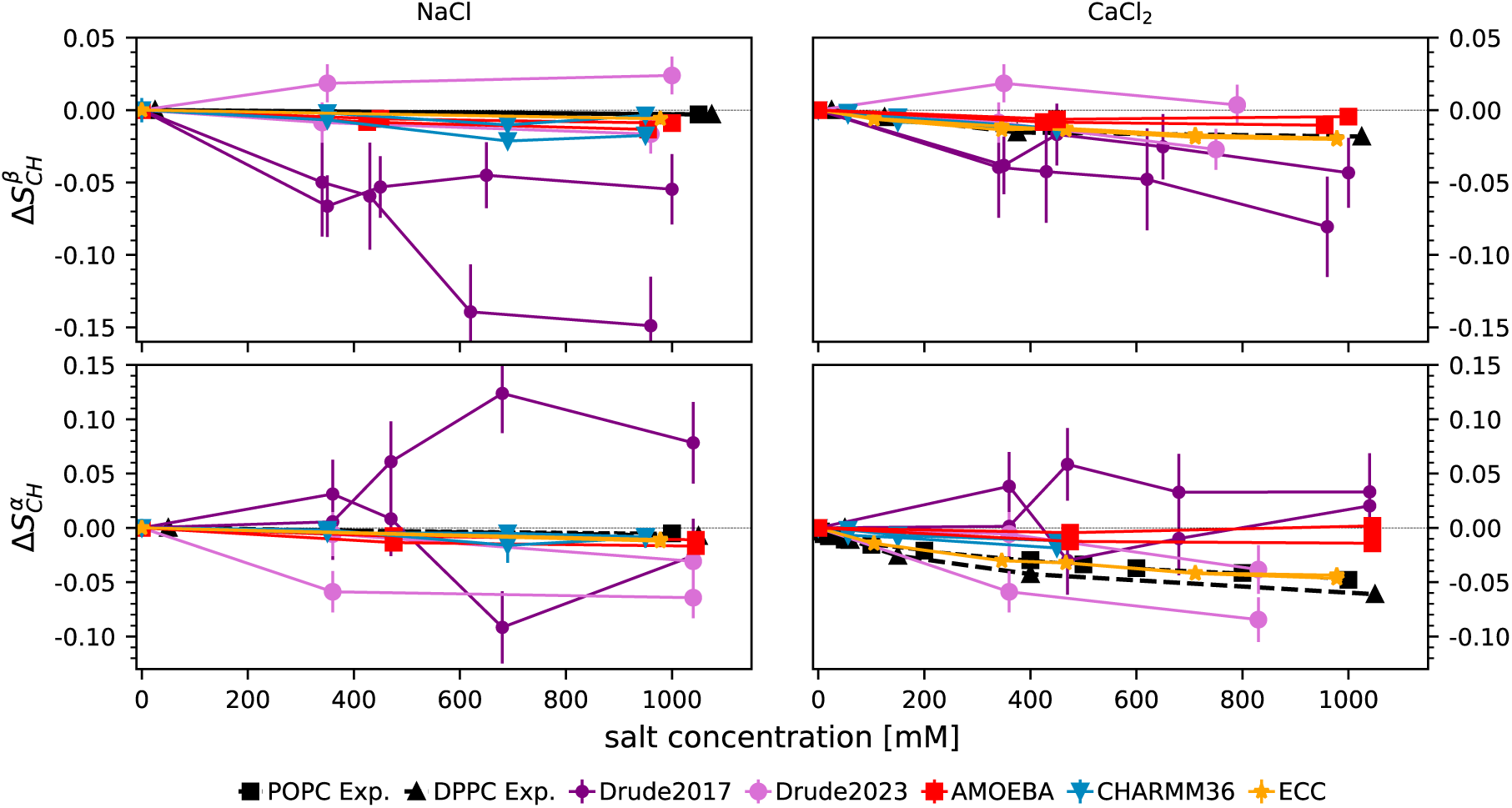
The change in the lipid head group order parameters *β* (top row) and *α* (bottom row) upon increasing ion concentration with respect to the simulations without salt. Data plotted separately for the two hydrogens attached to each carbon. CHARMM36 and EC-Clipids data are reproduced using the Zenodo repositories at Refs. 123–126 and Ref. 127, respectively. Note that the *y*-axis is trimmed to focus on the experimental data.^128,129^ A zoomed-in version of this figure is given in SI Fig. S4.

**Figure 4:**
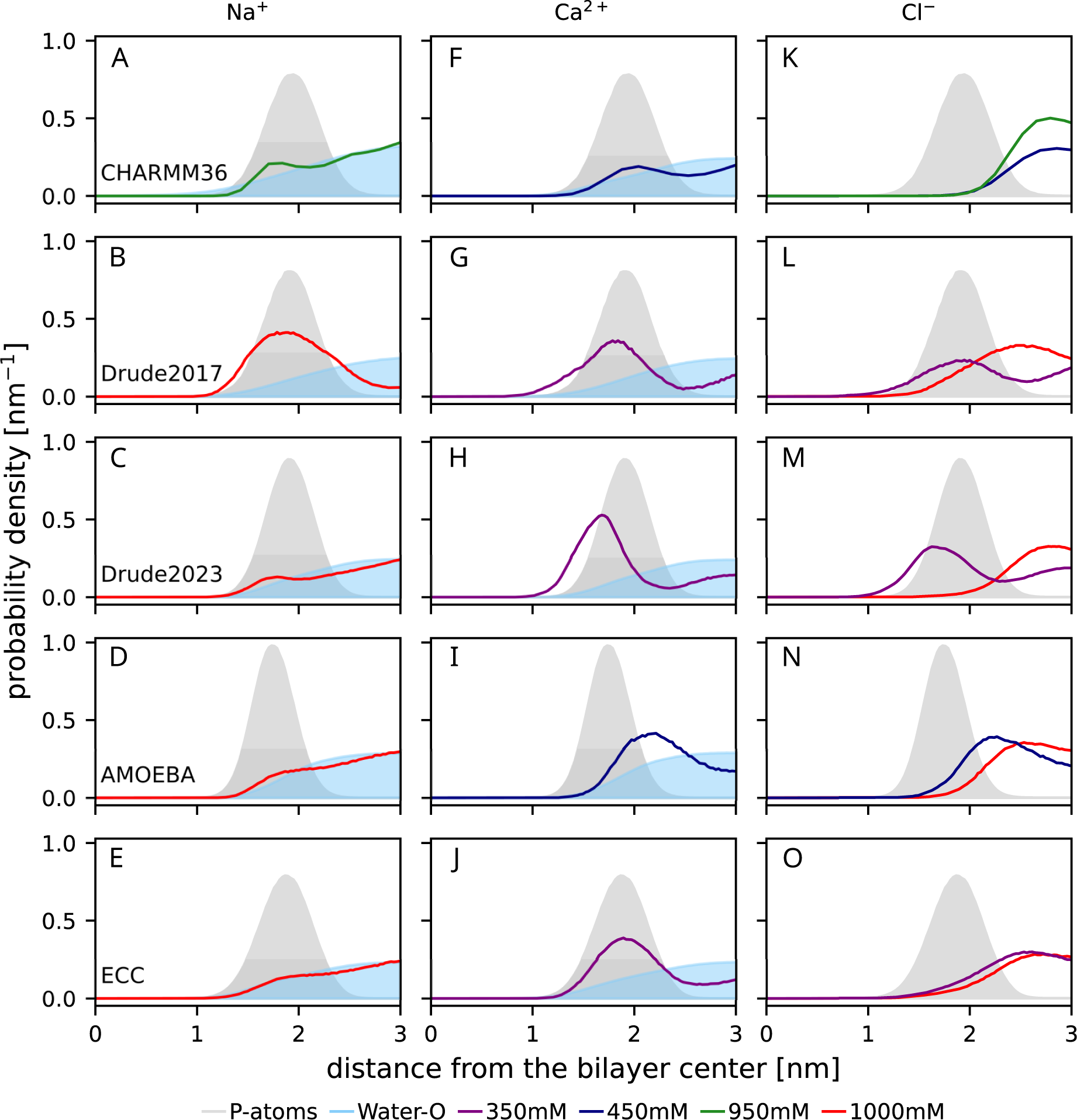
Density profiles along the membrane normal for (from the top): CHARMM36, CHARMM-Drude2017, CHARMM-Drude2023, AMOEBA, and ECClipids. In the third column, the Cl*^−^* densities are shown in the same color as their cations in the first and second columns. Note that for CaCl_2_, 350 mM (Drude models and ECC) and 450 mM (AMOEBA and CHARMM36) concentrations are shown; while for NaCl, 1000 mM concentration is shown for all force fields, except CHARMM36 (950 mM NaCl). The CHARMM36 data are reproduced using the Zenodo repositories of Refs. 123–126, ECC using the Zenodo repository of Ref. 127. Data are from POPC simulations for all force fields other than AMOEBA (DOPC).

CHARMM-Drude2017 predicts (Fig. 4G) similar calcium ion density profile as the model that is in good agreement with experiments (ECClipids, Fig. 4J). However, the sodium binding is equally strong in the CHARMM-Drude2017 simulations (Fig. 4B)—in contrast with both the ECC (Fig. 4E) and the experimental evidence: ^33^ The response of the headgroup order parameters to bound ions is not in qualitative agreement with experiments (Fig. 3). In particular, increasing 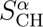 and the detectably different response of the two C–H bonds are not observed in experiments. This is in contrast with the results from previous bechmarking of non-polarizable simulations, ^33^ where the experimentally observed decrease of 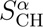 and 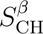 to more negative values upon ion binding were observed to be produced by all simulations (see Fig. 3 of Ref. 33), even though the binding affinity was often inaccurately predicted. This qualitative discrepancy in CHARMM-Drude2017 simulations may result from the incorrect lipid headgroup conformational ensemble (see Section 3.1), which leads to inaccuracies in the structural response of the ensemble to ion binding. Excessive sodium binding in the CHARMM-Drude model has been observed before for systems containing peptides or amino acids^116,130^ as well as deep-eutectic solvents.^131^

In simulations with the CHARMM-Drude2023 parameters, sodium and calcium ion binding are in line with the ECClipids simulations when comparing the cation density profiles (Fig. 4C,E;H,J)—although the calcium binding affinity is slightly larger, and Ca^2+^ ions penetrate deeper into the bilayer (Fig. 4H,J). The distributions of Cl*^−^* from CaCl_2_ show the largest difference: Whereas in ECClipids the Cl*^−^* density follows the water profile (Fig. 4O,J), in CHARMM-Drude2023 chloride penetrates deeper into the bilayer and echoes the Ca^2+^ profile (Fig. 4M,H). Interestingly, the latter feature is observed in all simulations with CaCl_2_ using polarizable force fields. However, sodium or calcium ion binding seem to again induce a response of different magnitude in the two C–H bonds attached to same carbon in CHARMM-Drude2023 simulations (Fig. 3) in contrast to experiments. This might indicate inaccurate structural response to ion binding, but poor convergence of the simulations owing to the slow conformational dynamics (see Section 3.2) cannot be ruled out in this case or in the case of the older 2017 version.

In the non-polarizable counterpart, CHARMM36 with the NBFIX correction, sodium binding (Fig. 4A) is similar to ECClipids (Fig. 4E) and CHARMM-Drude2023 (Fig. 4C), but the accumulation of anions outside the phosphate region is stronger (Fig. 4K,M,O). The structural response is well in line with the experiments: Similar to ECClipids but weaker than in CHARMM-Drude2023 (Fig. 3 left column). The divalent cation binding for CHARMM36 (Fig. 4F) is weaker than in ECClipids (Fig. 4J), and the *α* carbon order parameter response is smaller than in experiments and in ECClipids (Fig. 3 bottom right). Comparing the calcium distribution from CHARMM36 (Fig. 4F, which is rather similar to the Na^+^ distribution in Fig. 4A) with CHARMM-Drude2023 (Fig. 4H) suggests that the polarizable model may better capture the difference in the relative amounts of Na^+^ and Ca^2+^ bound than correcting the ion binding by scaling the Lennard-Jones parameters (NBFIX). That said, the overall structural response to ion binding in CHARMM36 appears more realistic (Fig. 3).

In the AMOEBA-based simulations, sodium binds weakly (Fig. 4D) and does not affect the order parameters (Fig. 3 left column), consistent with experiments and the ECClipids simulations (Fig. 4E). Calcium binding affinity (Fig. 4I) is similar to ECClipids (Fig. 4J), but the order parameters do not change upon binding contrasting the experiments (Fig. 3 right column). This may result from the binding position of calcium, which is outside the phosphate density peak in the AMOEBA simulations (Fig. 4I). In other simulations calcium penetrates to phosphate region or deeper and, in agreement with the “lipid electrometer”, ^120^ reorients the headgroup dipole, giving rise to changes in order parameters in line with the experimental data.^33^

In conclusion, incorporating explicit polarizability as implemented in the CHARMM-Drude or AMOEBA-based parameters used here does not necessarily lead to an improved description of cation binding to phospholipid membranes. These force fields do not correctly capture the response of lipid headgroup to cation binding, most likely due to inaccuracies in lipid parameters. As such inaccuracies can also affect ion binding, it is difficult to isolate the explicit influence of polarizability per se on ion binding.

### 3.4 Conclusions

Including electronic polarizability to MD simulation models of membranes is expected to improve the description of bilayer polar regions and their interactions with charged molecules, thereby making MD simulations of complex biomolecular systems more realistic. However, quality of polarizable membrane simulations has not been evaluated on an equal footing with the non-polarizable ones. Here, we used the quality evaluation metrics defined in the NMRlipids Databank^40^ together with additional analyses on dynamics and ion binding to evaluate the performance of two available polarizable lipid model types, the CHARMM-Drude^19,26^ and the AMOEBA-based^20,27^ parameters, against experimental NMR and SAXS data and the best-performing non-polarizable force fields. Considering the complexity and additional computational cost of simulations with polarizable models, it is crucial to understand their accuracy with respect to experiments and to choose the best models according to their respective strengths when planning simulations.

Our comparisons of lipid conformations and dynamics show that there is room for improvement in the current polarizable parameterizations, even to reach the level of the best currently available non-polarizable force fields. Although the most recent CHARMM-Drude23 has improved the description of molecular conformations and dynamics, both tested CHARMM-Drude models predict a slightly too ordered membrane core and vastly too slow headgroup dynamics. The latter can compromise the convergence of the simulations within the typically used simulation times. This notion is further supported by the large relative equilibration times detected for the CHARMM-Drude models. The tested AMOEBA-based parameters have difficulties capturing ordering in the lipid acyl chain region and other more general membrane properties interconnected with chain conformations; yet, the description of headgroup conformations is relatively good, and the dynamics have similar quality as in the best non-polarizable force fields.

Sodium and calcium binding to membranes in simulations were evaluated using the experimentally observed headgroup C–H bond order parameter changes upon addition of NaCl or CaCl_2_. The binding in explicitly polarizable models was compared with the ECClipids model, which implicitly includes electronic polarizability and gives the currently most accurate response to ion binding, and with the non-polarizable CHARMM36, which when used with the NBFIX ion model also rivals its polarizable counterparts. Compared to CHARMM36, CHARMM-Drude2023 provides an improved description of the stronger binding of calcium compared to sodium; this difference between sodium and calcium binding is also present in simulations with the AMOEBA-based parameters. However, the calcium binding depth, affinity, and the consequent structural response of the lipids do not exactly align with the experiments (or with the ECClipids results) either in CHARMM-Drude2023 or AMOEBA. The incorrect response to ion binding likely connects to the other discussed inaccuracies in lipid conformational ensembles, dynamics, and membrane order.

In summary, the potential and promise of explicitly polarizable lipid force fields to improve the description of bilayer membranes have not yet been fully realized. However, it seems likely that this is not an inherent flaw in polarizability, but rather in the current parameterizations and their incompatibility with parameters describing other interactions, such as those for Lennard-Jones interactions. As the non-polarizable force fields benefit from a longer history of development and more intense scrutiny, it is not surprising that they currently out-perform their polarizable counterparts. On the other hand, the marked improvements from CHARMM-Drude2017 to CHARMM-Drude2023 demonstrate the potential of parameter tuning in improving polarizable force fields. Such endeavors are expected to substantially ease with the emerging automated methods for parameter development.^26,37^

## Author contributions

H.S.A. analyzed the correlation and relaxation times from the simulations, coordinated the project, and participated in the writing of the manuscript from the first version onwards. S.D. assisted in generating the initial system setups for the simulations of POPC and POPE bilayers using the AMOEBA-based force field (including identifying errors in the force field files received), and wrote the initial scripts required to generate OpenMM-and-Tinker-suitable file formats of the simulation systems. B.K. conceptualized and initiated the project, did the literature research, created and ran the simulations, analyzed the results, and wrote the manuscript. J.J.M. contributed to early testing (debugging, setup, and running) of simulations with the AMOEBA-based force field using Tinker9 and Tinker-OpenMM as well as manuscript editing. M.S.M. assisted in conceptualizing the project and critically refined the manuscript. O.H.S.O. made the quality evaluation using the NMRlipids Databank, and assisted in conceptualizing the project and in writing the manuscript.

## Acknowledgement

All authors collectively would like to thank to Dr. Venable for making their CHARMM-Drude2023 simulation trajectories publicly available. B.K. thanks Huiying Chu and Guohou Li for providing the AMOEBA-based force field parameters and technical discussions, and Gianni Klesse for providing the AMOEBA-based force field parameters in OpenMM format and technical help with running the simulations. J.J.M. thanks Huiying Chu (from the Li lab) for technical discussions and Sameer Varma for useful suggestions. M.S.M. acknowledges support by the Trond Mohn Foundation (BFS2017TMT01). O.H.S.O. acknowledges CSC – IT Center for Science for computational resources and the Research Council of Finland for funding (grant nos. 315596, 319902, 345631 & 356568).

## Supporting Information Available

Supporting Information: NMR order parameters and SAXS form factors for additional CHARMM-Drude2023 replicas, dihedral angle distributions for the headgroup from the studied polarizable models and CHARMM36, extended version of Fig. 3, and the bilayer electron density profiles for the studied polarizable models. This information is available free of charge via the Internet at http://pubs.acs.org

