## Supplementary Information for "Evaluating polarizable biomembrane simulations against experiments"

### Supplementary Information: Evaluation of polarizable biomembrane simulations against experiments

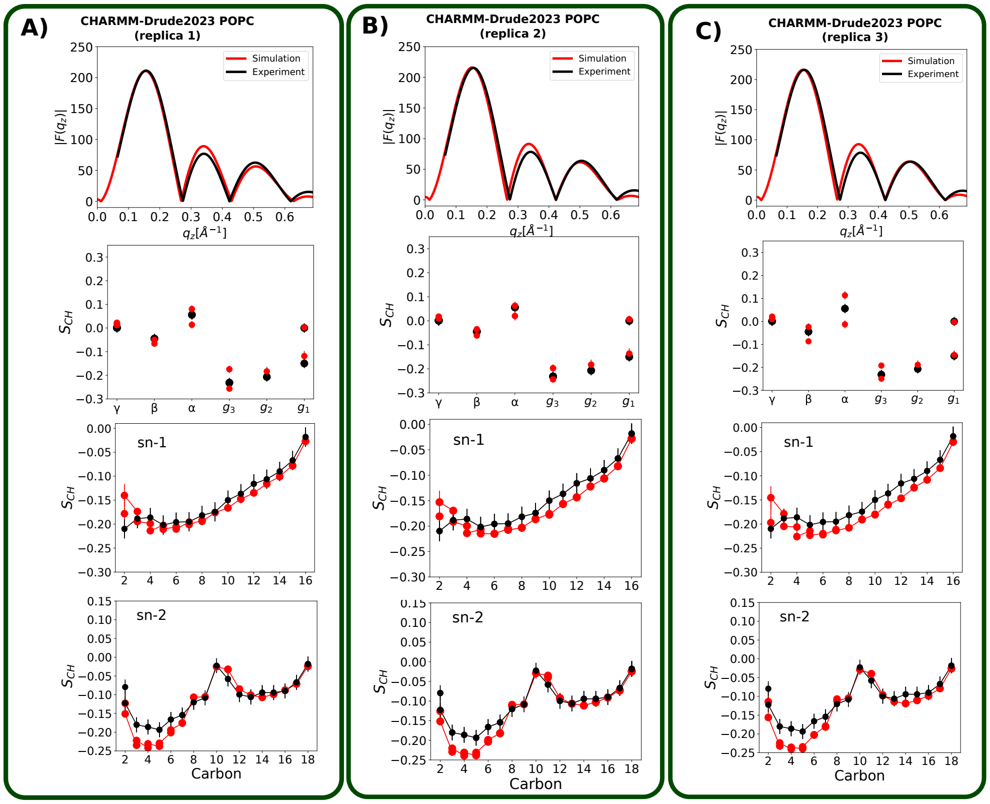


Figure S1: Comparison of different replicas from CHARMM-Drude2023 simulations against experimental X-ray scattering form factors and C–H bond order parameters from the NMRlipids Databank.

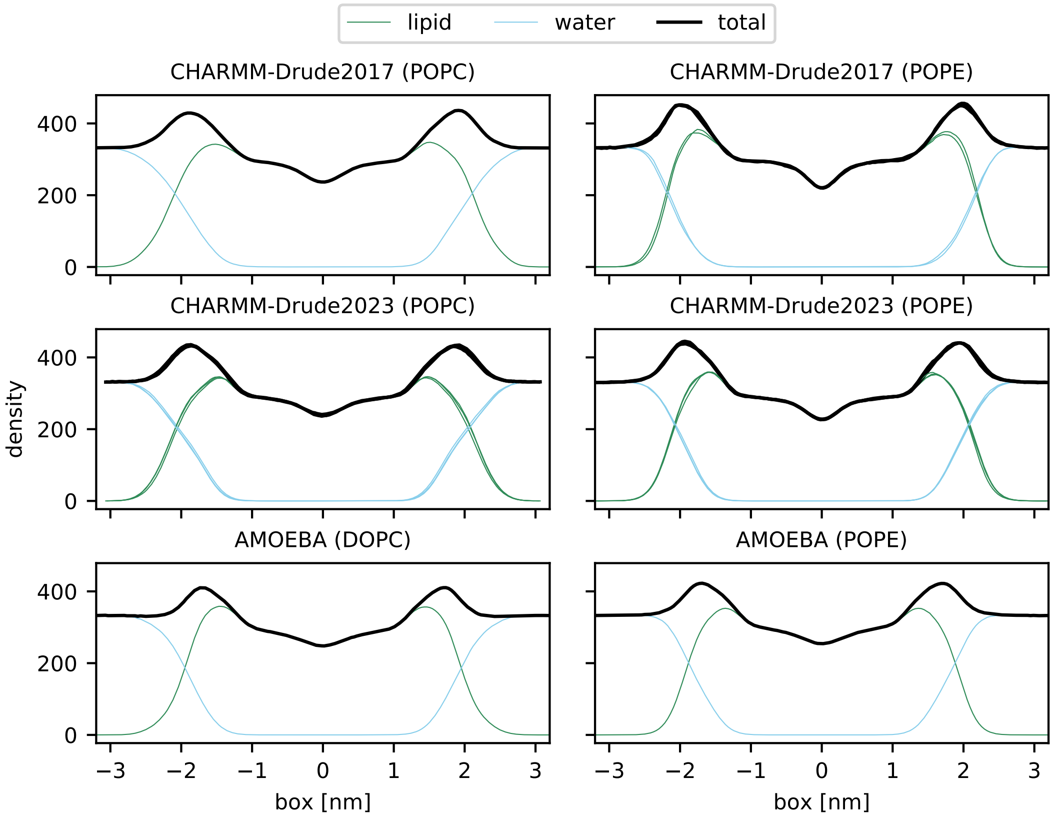


Figure S2: The bilayer electron density profiles from salt-free simulations with the investigated polarizable models. The data are centered such that the bilayer center resides at zero. For some systems, multiple replicas are plotted as indicated by the presence of multiple (mostly overlapping) lines.

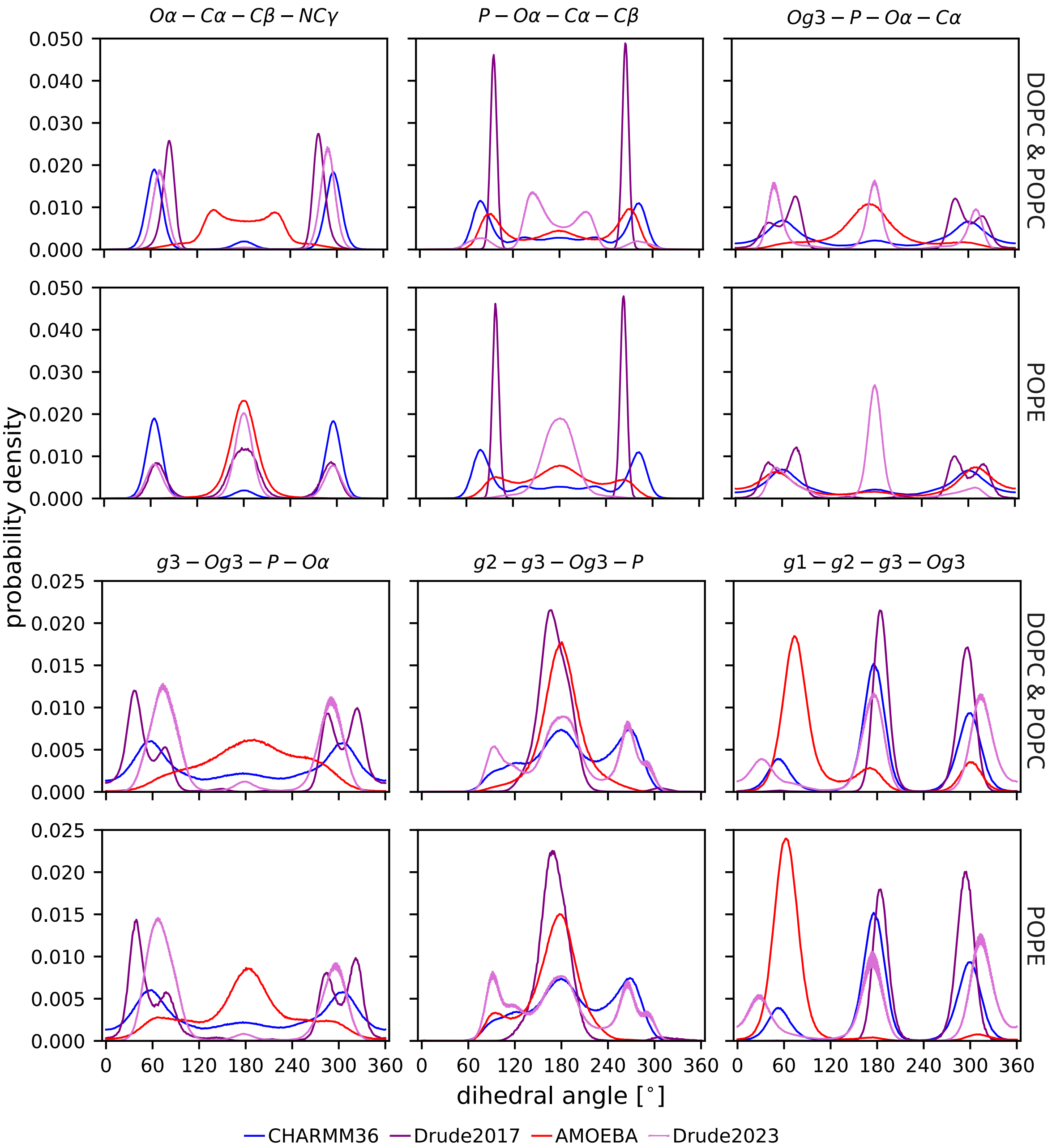


Figure S3: Distributions of the torsion angles for head group atoms. The atoms forming the dihedral angles are given in the titles. For each torsion angle, the upper rows contain DOPC (AMOEBA) and POPC (CHARMM-Drude2017/CHARMM-Drude2023) data and the lower rows contain dihedrals for POPE.

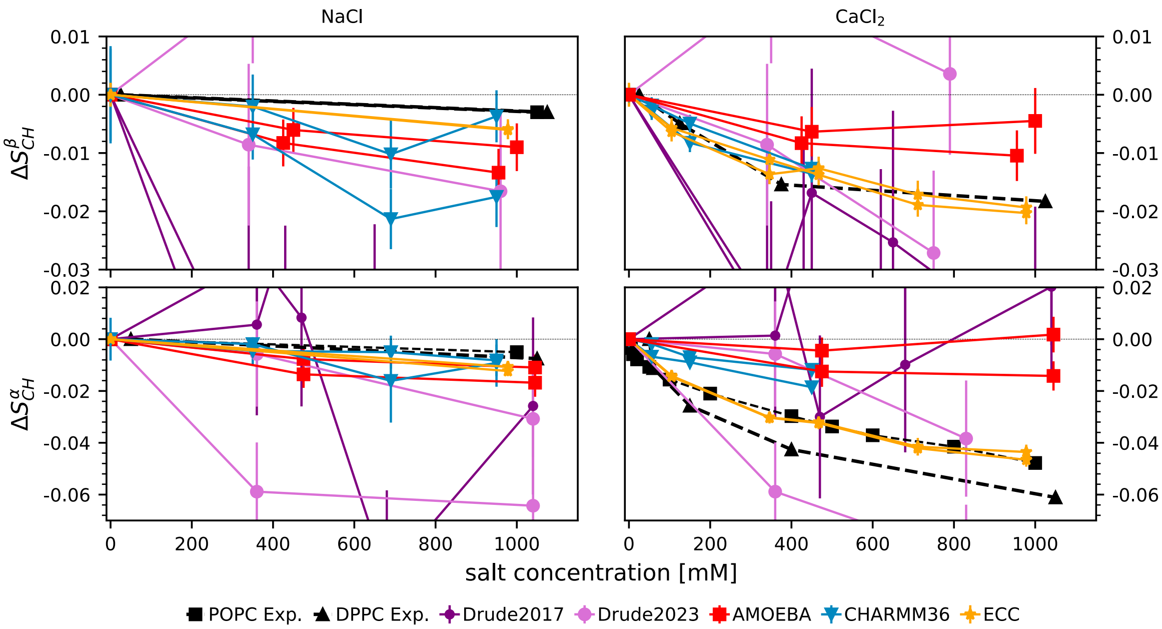


Figure S4: The change in the lipid head group order parameters $\beta$ (top row) and $\alpha$ (bottom row) upon increasing ion concentration with respect to the simulations without salt. Data plotted separately for the two hydrogens attached to each carbon. CHARMM36 and ECClipids data are reproduced using the Zenodo repositories at Refs. 1–4 and Ref. 5, respectively.
